# Synaptic vesicles dynamics in neocortical epilepsy

**DOI:** 10.1101/2020.01.05.895029

**Authors:** Eleonora Vannini, Laura Restani, Marialaura Dilillo, Liam McDonnell, Matteo Caleo, Vincenzo Marra

## Abstract

Neuronal hyperexcitability often results from an unbalance between excitatory and inhibitory neurotransmission, but the synaptic alterations leading to enhanced seizure propensity are only partly understood. Taking advantage of a mouse model of neocortical epilepsy, we used a combination of photoconversion and electron microscopy to assess changes in synaptic vesicles pools *in vivo*. Our analyses reveal that epileptic networks show an early onset lengthening of active zones at inhibitory synapses, together with a delayed spatial reorganization of recycled vesicles at excitatory synapses. Proteomics of synaptic content indicate that specific proteins were increased in epileptic mice. Altogether, our data reveal a complex landscape of nanoscale changes affecting the epileptic synaptic release machinery. In particular, our findings show that an altered positioning of release-competent vesicles represent a novel signature of epileptic networks.

## Introduction

Epilepsy is a disorder of the central nervous system that affects around 50 million people worldwide and it is characterized by recurrent spontaneous seizures, that are the clinical manifestation of an excessive hypersynchronous discharge of a population of neurons (Bromfield and Cavazos, 2006). Despite all the available treatments, one third of patient develop a drug-resistant form of epilepsy (Kwan et al.; Sharma et al., 2015). The propensity to develop seizures is due to brain damage, in lesional epilepsy (Pitkänen and Immonen, 2014; Wasilewski et al., 2020; Wie Børsheim et al., 2020), or to an altered synaptic function at excitatory and/or inhibitory terminals, in non-lesional epilepsy (Wykes et al., 2012; Farisello et al., 2013; Corradini et al., 2014; Ferecskó et al., 2015; Snowball et al., 2019). Specifically, chronic epilepsy is thought to result from a synaptic reorganization that leaves permanent marks on cortical networks and may lead to network dysfunction, cognitive deficits and impaired information processing (Pitkänen et al., 2013, 2015; Holmes, 2015; Vannini et al., 2016). Epileptic networks display several modifications in synaptic function and structure at the level of presynaptic boutons, post-synaptic structures and the glial processes enwrapping them (Bernard, 2010). In the presynaptic compartment, recent technological advances have allowed a detailed characterization of the size and spatial organization of functional vesicle pools. These parameters correlate with measures of synaptic strength and are altered following plasticity-inducing stimuli (Rey et al., 2020). However, such ultrastructural readouts of synaptic function have not been applied thus far to the study of epileptogenic modifications or their consequences.

The correct functioning of neuronal networks requires precise modulation of excitatory and inhibitory activity (Xue et al., 2014; Rossi et al., 2017; Rubin et al., 2017; Sohal and Rubenstein, 2019). When network activity is tipped out of balance, a number of cellular processes take place to re-establish its normal function (Turrigiano, 2012). The processes underlying homeostatic plasticity can affect cellular activity and synaptic output (Wefelmeyer et al., 2016). Following a perturbation, neurons attempt to restore their baseline firing rates and dynamic range by regulating their intrinsic excitability, probability of neurotransmitter release and neurotransmitter receptor expression (Davis and Müller, 2015). Central synapses have the ability to plastically adapt to new conditions by dynamically scaling up and down their output. Such regulation has been demonstrated to occur during development, sleep and learning. However, much less is known about the mechanisms of homeostatic scaling as a consequence of a pathological, system-level perturbation *in vivo* (Turrigiano, 2008, 2012; González et al., 2019).

Here we took advantage of a well-characterised model of chronic, focal epilepsy in the visual cortex (Mainardi et al., 2012; Chang et al., 2018) to investigate synaptic changes in hyperexcitable networks. Tetanus neurotoxin (TeNT) is a metalloprotease that cleaves the synaptic protein VAMP/synaptobrevin leading to the establishment of a focal cortical hyperexcitability, with electrographic seizures that persist for several weeks after TeNT wash out (Nilsen et al., 2005; Jiruska et al., 2010; Mainardi et al., 2012; Vannini et al., 2016; Snowball et al., 2019). Despite many studies characterizing TeNT-induced epilepsy (i.e. seizures manifestation), very little is known about the persistent synaptic changes that underpin chronic, spontaneous seizures. The absence of neuronal loss and gliosis, together with the persistence of spontaneous seizures, chronically altered neural processing and structural modifications of both dendritic spines and branches suggest that synaptic modifications occur and last after TeNT clearance (Mainardi et al., 2012; Vannini et al., 2016). Here, using FM1-43FX and activity-dependent labelling of synaptic vesicles, we simultaneously investigated function and ultrastructure of both excitatory and inhibitory terminals, in acute and chronic phases of TeNT-induced epilepsy. We combined this approach with an unbiased measure of proteins content in the two phases of hyperexcitability, isolating synaptosomes at different time points after TeNT injection, and assessing by electrophysiological recordings the impact of inhibiting Carboxypeptidase E (CPE), upregulated in epileptic mice, on seizures occurrence.

## Methods

### Animals and TeNT injections

Adult (age > postnatal day 60) C57BL/6J mice used in this study were reared in a 12h light-dark cycle, with food and water available *ad libitum*. All experimental procedures were conducted in conformity with the European Communities Council Directive *n*° 86/609/EEC and were approved by the Italian Ministry of Health. TeNT (Lubio; Lucerne, Switzerland; 0.1—0.2 ng) or RSA (Rat Serum Albumin) solutions in PBS were intracranially injected into the primary visual cortex (i.e. 0.0 mm anteroposterior, 2.7 mm lateral to the lambda suture and at a cortical depth of 0.65 mm) ; of anesthetised (ketamin/Xylazine 100-10 mg/Kg) mice. After surgery, a glucose solution (5 % in saline) was subcutaneously administered and recovery of animals was carefully monitored. Paracetamol was added in drinking water for 3 days. Additional details can be found in Mainardi et al., 2012; Vallone et al., 2016; Vannini et al., 2016, 2017. No behavioural seizures are detectable in those animals, as already reported in (Mainardi et al., 2012; Vannini et al., 2016).

### FM 1-43FX injection and visual stimulation

Control and epileptic mice, deeply anesthetized with urethane (7 ml/kg; 20 % solution in saline, i.p.; Sigma) and placed in a stereotaxic apparatus, received an injection of FM 1-43FX dye into the primary visual cortex, layers II-III (i.e. 0.0 mm anteroposterior and 2.7 mm lateral to the lambda suture, 0.7 mm depth). FM1-43FX is a non-permeable styryl dye that labels the cell membrane. Once synaptic vesicles fuse with the cell membrane, molecules of FM1-43FX diffuses laterally along the membrane previously comprising the vesicle. As vesicles undergo endocytosis, as part of their recycling, the dye is trapped inside the recently released vesicles (Marra et al., 2014). 3 min later, animals were stimulated for 10 min with square-wave gratings (1 Hz, 0.06 c/deg, contrast 90%) and flashes of light. All visual stimuli were computer-generated on a display (Sony; 40 × 30 cm; mean luminance 15 cd/m2) by a VSG card (Cambridge Research Systems). Mice, still under anaesthesia, were kept in the dark and perfused through the heart with a fresh solution of 6% glutaraldehyde, 2% formaldehyde in PBS, as described in (Jensen and Harris, 1989) right after the end of the visual stimulation

### Photoconversion and Electron Microscopy analysis

All the following procedures were made in the dark. The protocol followed is described in detail in (Marra et al., 2014). Briefly, embedded in EPON, slices were collected with an ultramicrotome serial sections (70 nm thickness) and placed in grids at RT. Thereafter, sections could be viewed with a transmission electron microscope fitted with a cooled CCD camera. Images were acquired using local landmarks to identify the same target synapse in consecutive sections and analysed using Image J/Fiji (NIH) and a custom script in Python (Python.org). At ultrastructural level, target synapses (visual cortex, layers II-III) were randomly chosen and synaptic vesicles were scored based on their vesicle luminal intensity using methods outlined previously (Darcy et al., 2006), image names were changed to ensure that the experimenters were blind to the experimental condition of each electron micrograph. A terminal was considered inhibitory if no spine or postsynaptic density could be observed in the middle section and in at least one of the adjacent sections. As expected (Meyer et al., 2011; Tremblay et al., 2016; van Versendaal and Levelt, 2016; Lim et al., 2018), inhibitory terminals were estimated to be 15-25% of the total.

### Synaptosomes extraction and proteomic analysis

Synaptosomes were extracted using a slightly modified protocol taken from (Giordano et al., 2018). Visual cortices were gently homogenized in 500 μl of ice cold homogenizing buffer (0.32 M sucrose, 1 mM EDTA, 1mg/ml BSA, 5 mM HEPES pH 7.4, proteases inhibitors) and centrifuged 10 min at 3000 g at 4°C; supernatant was recovered and centrifuged again for 15 min at 14000 g at 4°C. After discarding supernatant, the pelleted synaptosomes were suspended in 110 μl of Krebs-Ringer Buffer and 90 μl of Percoll (Sigma-Aldrich) were added. A 2 min spin (14000 rpm, 4°C) was performed and enriched synaptosomes were recovered from the surface of the solution with a P1000 tip and resuspended in 1ml of Krebs-Ringer buffer. After an additional spin of 2 min (14000 rpm, 4°C), the supernatant was discarded and the pellet resuspended in 20 μl of RIPA buffer.

### Proteomics sample preparation and data analysis

Trypsin/LysC mix Mass Spec grade was purchased from Promega (Madison, WI). Tandem Mass Tags (TMT 10-plex) kits and microBCA protein assay kit were purchased from Thermo Fisher Scientific (Rockford, IL). All other reagents and solvents were purchased from Sigma-Aldrich (St. Louis, MO). Synaptosomes proteome extracts were quantified with a micro BCA protein assay and aliquots of 3.5 μg of proteins were diluted to 40 μL of RIPA/Trifluoroethanol (TFE) 50/50. Paramagnetic beads were added to each sample and further processed following a modified SP3 protocol for ultrasensitive proteomics as previously described (Pellegrini et al., 2019). Synaptosomes proteins were reduced alkylated and digested with a mixture of trypsin/Lys-C (1:20 enzyme to protein ratio). Digested peptides were then quantified, and labelled with TMT 10-plex: samples were block randomized (www.sealedenvelope.com) over 5 TMT sets. Each TMT set included 2 normalization channels for batch corrections built pooling an aliquot from each digested synaptosome sample (Plubell et al., 2017). TMT sets underwent high pH fractionation on an AssayMap Bravo (Agilent technologies) and fractions run on a nano-LC (Easy1000 Thermo Fisher Scientific) equipped with a 50 cm EasySpray column and coupled with an Orbitrap Fusion for MS3 analysis (Thermo Fisher Scientific). Experimental details regarding sample fractionation and LC-MS/MS runs have been already reported elsewere (Pellegrini et al., 2019). Data were analysed using Proteome Discoverer 2.1. TMT data were normalized by internal reference scaling (Plubell et al., 2017).

### Electrophysiological recordings and drugs administration

Surgery was performed as described in (Spalletti et al., 2017), but the small craniotomy was centred at 3 mm lateral to Lambda and performed in TeNT/RSA-injected hemisphere. Neuronal activity was recorded with a NeuroNexus Technologies 16-channel silicon probe with a single-shank (A1×16-3mm-50-177) mounted on a three-axis motorized micromanipulator and slowly lowered into the portion of visual cortex previously injected with TeNT or RSA solution. The tip of the probe was positioned at the depth of 1 mm so that the electrode contacts (spaced by 50 microns) sampled activity from all cortical layers. Before the beginning of the recording, the electrode was allowed to settle for about 10 min. Local Field Potentials (LFP) signals were acquired at 1 kHz and bandpass filtered (0.3 Hz to 200 Hz) with a 16-channel Omniplex recording system (Plexon, Dallas, TX). Local Field Potentials (LFP) were computed online and referred to the ground electrode in the cerebellum. In order to verify whether interacting with Carboxypeptidase E (CPE) would change epileptic activity, we topically applied over the craniotomy 2 μL of PBS containing 25 μM of 2-guanidinoethylmercaptosuccinic acid (GEMSA; Sigma-Aldrich) without removing the electrode. Neural signals were acquired at regular time intervals up to 30 min after GEMSA delivery to verify the effect and the penetration of the drug in the cortical layers. At the end of the experiment animals were sacrificed. Data were analysed offline with NeuroExplorer software (Plexon Inc, USA) and with custom made Python interfaces (Python.org). Movement artifacts were removed offline. The coastline analysis was calculated as the sum of the absolute difference between successive points (Wykes et al., 2012).

### Statistical analysis

Statistical analysis was performed with Graph Pad (version 8) except for proteomics analysis, in which we used Perseus. Normality of distributions was assessed with D’Agostino test and appropriate test was chosen accordingly.

## Results

### Ultrastructural investigation of synaptic vesicle function in hyperexcitable networks

Our studies included three groups of C57BL/6 mice with injections into the primary visual cortex (V1): a control group injected with vehicle (Control), an Acute epileptic group tested 10 days after TeNT injection in V1 and a Chronic epileptic group tested 45 days after TeNT injection in V1. Synaptic vesicles from 3 animals in each of these groups were labelled by infusing FM1-43FX in the visual cortex while presenting a series of visual stimuli (**Fig. 1A**). After fixation, photoconversion and processing for electron microscopy, we were able to label individual vesicles at excitatory (asymmetrical) and inhibitory (symmetrical) synapses (**Fig. 1A**). This approach allowed us to identify individual synaptic vesicles, released and recycled in the presence of FM1-43FX, as having an electron dense lumen, while non-released vesicles present a clear lumen and a darker membrane (Marra et al., 2012). First, we investigated the length of the active zone as a readout of synaptic activity independent of our labelling protocol (Harris and Weinberg, 2012). Surprisingly, we found an increase in the length of inhibitory synapses’ active zone in the acute phase (**Fig. 1B**), normally associated with increased release. However, the released fraction of inhibitory vesicles (number of released vesicles over total number of vesicles) is reduced in animals injected with TeNT (**Fig. 1C**), shown to preferentially impair inhibitory release (Schiavo et al., 2000). No differences in active zone length and released fraction of vesicles were found at excitatory synapses (**Fig. 1B, 1C**). Interestingly, the total number of vesicles across control and epileptic groups does not change significantly neither at excitatory (Control: mean= 52.16, SD=35.47, n= 49; Acute: mean= 56.23, SD=29.81, n=59; Chronic: mean= 51.28, SD=19.39, n= 124; data not shown) nor inhibitory synapses (Control: mean= 40.76, SD=14.38, n=17; Acute: mean= 54.33, SD=36.12, n=18; Chronic: mean= 54.33, SD=19.91, n=30; data not shown).

**Figure 1:**
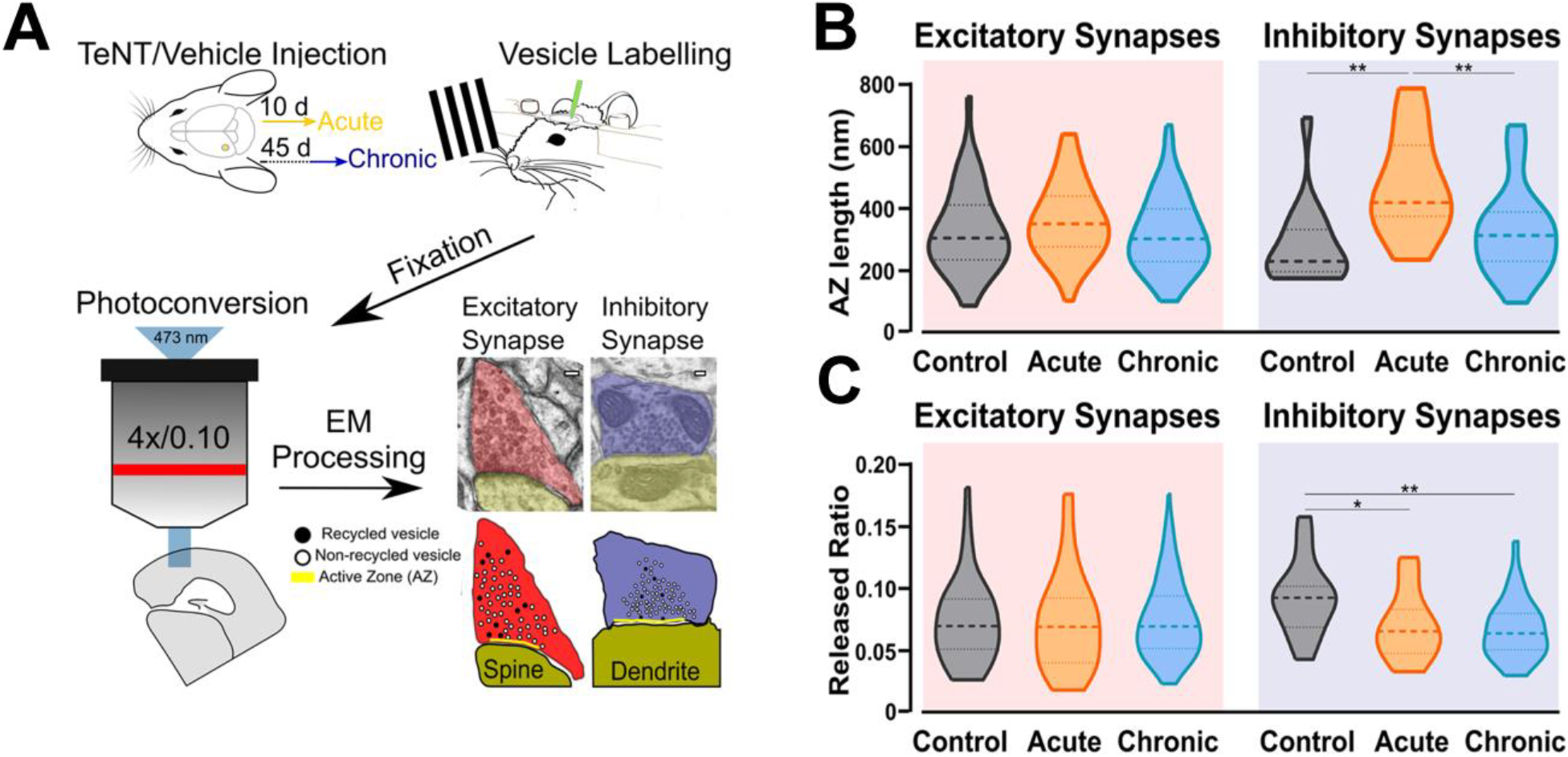
Ultrastructural and functional changes at presynaptic terminals of TeNT-injected mice. **A**) Diagrammatic representation of labelling protocol. Visual cortices from mice in Control, Acute and Chronic groups were infused with FM1-43FX during mild visual stimulation. Brains were rapidly fixed and sliced to allow photoconversion of FM1-43FX signal before processing for electron microscopy. Individual presynaptic terminals were classified as excitatory (asymmetrical synapses, red) or inhibitory (symmetrical synapses, blue); size, position and numbers of active zone (AZ, yellow), non-released vesicles (open circles) and released vesicles (black circles) were analysed (scale bars 100 nm). **B**) Left: Active zone (AZ) length in Control (grey), Acute (orange) and Chronic (blue) epileptic mice at excitatory synapses; no differences between the groups (Kruskal-Wallis test, p = 0.10). Distribution, median and quartiles shown for each group; Control n=41; Acute n=46; Chronic n=118. Right: Active zone (AZ) length in Control (grey), Acute (orange) and Chronic (blue) epileptic mice at inhibitory synapses (Kruskal-Wallis test, p < 0.01, Control vs Acute p<0.01, Control vs Chronic p>0.05, Chronic vs Acute p<0.01). Distribution, median and quartiles shown for each group; Control n=15; Acute n=14; Chronic n=29. **C)** Left: Released fraction of synaptic vesicles (labelled vesicles/total vesicles) in Control (grey), Acute (orange) and Chronic (blue) epileptic mice at excitatory synapses; no differences between the groups (Kruskal-Wallis test, p = 0.67). Distribution, median and quartiles shown for each group; Control n=47; Acute n=57; Chronic n=118. Right: Released fraction of synaptic vesicles of Control (grey), Acute (orange) and Chronic (blue) epileptic mice at inhibitory synapses. (Kruskal-Wallis test, p < 0.05, Control vs Acute p<0.05, Control vs Chronic p<0.01, Chronic vs Acute p>0.05). Distribution, median and quartiles shown for each group; Control n=16; Acute n=16; Chronic n=30.

### Changes in docking and positioning of activated vesicles at excitatory synapses in chronic epilepsy

After quantifying direct and indirect measures of vesicular release, we examined the spatial distribution of released and non-released vesicles within presynaptic terminals. We started by analysing the released fraction in the docked and non-docked populations of vesicles. As described before for excitatory synapses (Marra et al., 2012), the Control group showed a higher released fraction in the docked population, similar results were found in the Acute group. Conversely, in the Chronic group the released fraction was higher in the non-docked population at excitatory synapses (**Fig. 2A**). We also report that inhibitory synapses have a higher released fraction in the docked population, which does not seem to be affected by the induction of epilepsy (**Fig. 2A**). To gain insight on the effect observed at excitatory synapses of the Chronic group, we compared the distance of released and non-released vesicles from the active zone (**Fig. 2B**). We reasoned that if the effect is specific to the ability of released vesicles to dock, their position within the terminal should not be affected. We examined the cumulative fraction of the distance of released and non-released vesicles from their closest point on the active zone. At excitatory synapses, in Control and Acute groups the released vesicle population are closer to the active zone compared to the non-released population. However, in the Chronic group released excitatory vesicles do not show a spatial bias towards the active zone, that was instead observed in the other groups (**Fig. 2B**). At inhibitory synapses, the distances of vesicular populations to the active zone has a different pattern, with no difference between released and non-released population in Control and Acute groups and with a spatial bias of released vesicles towards the active zone in the Chronic group (**Fig. 2B**). As a visual representation of the distribution of released vesicles at excitatory and inhibitory across the three conditions, we generated 2D histograms of the distribution of released vesicles within spatially normalised terminals, with the centre of the active zone at the origin of the X axis (**Fig. 2C**). This representation shows a clear broadening of the distribution of released excitatory vesicles in the chronic phase.

**Figure 2:**
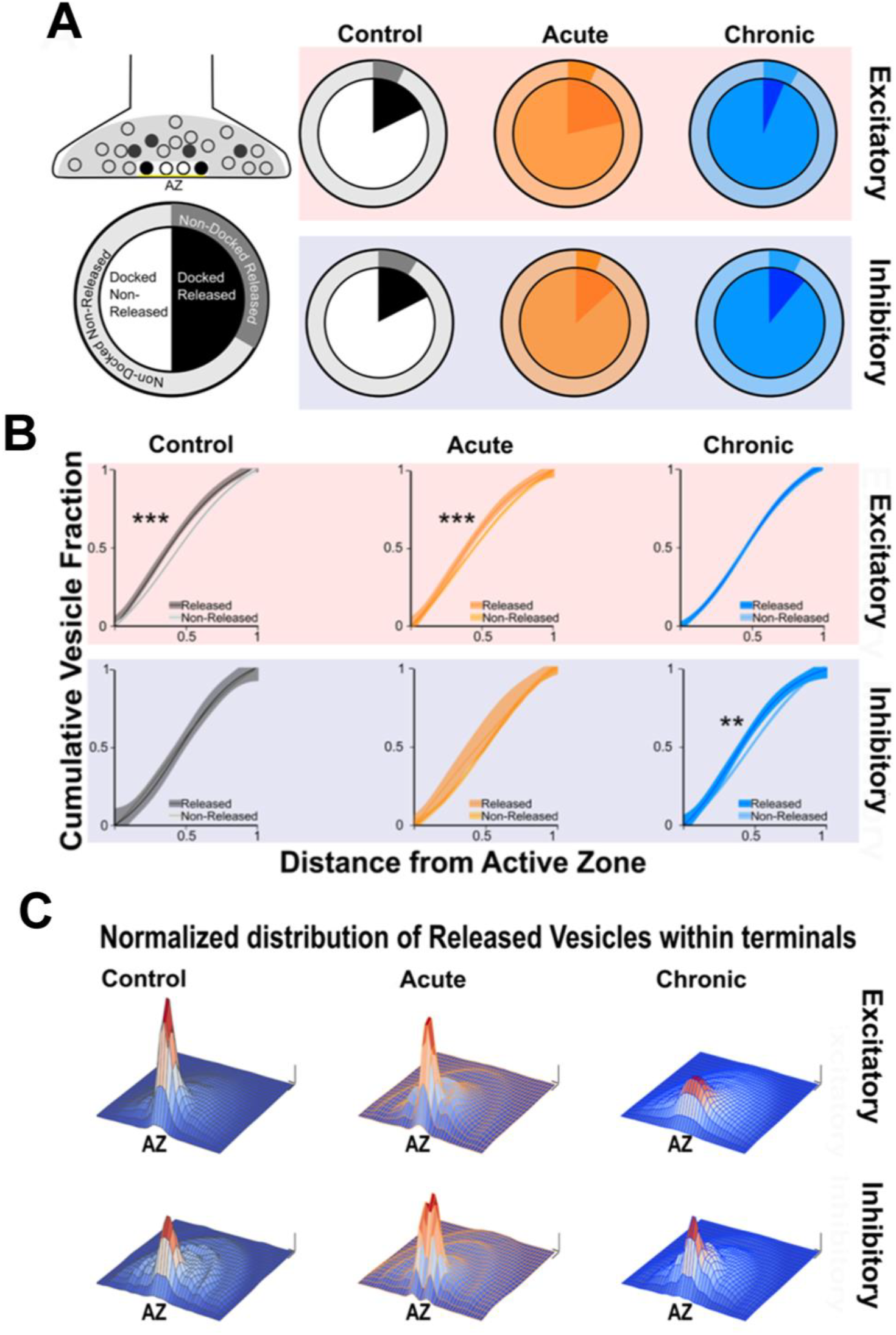
Changes in released vesicles’ docking and spatial organization in chronic phase of epilepsy. **A**) Ratio of released vesicles in the docked and undocked population. Left: Diagram and legend for each pie chart. Top: Excitatory synapses’ ratio of released vesicles (darker) in docked (inner pie chart) and undocked population (outer pie chart) in Control (grey), Acute (orange) and Chronic (blue) groups. Only the Chronic group shows a significant difference from expected frequencies based on control observation (Chi-squared test: p< 0.001). Bottom: Inhibitory synapses’ ratio of released vesicles (darker) in docked (inner pie chart) and undocked population (outer pie chart) in control (grey), acute (orange) and chronic (blue) groups. **B**) Distance of released or non-released vesicles to the closest point on the active zone. Left: Diagram representing of how distance measures were taken at each synapse. Top: Sigmoid fit and 95% confidence interval of cumulative fraction of distance between released and not-released synaptic vesicles to the active zone at excitatory synapses in Control (grey), Acute (orange) and Chronic (blue) epileptic mice. Bottom: Sigmoid fit and 95% confidence interval of cumulative fraction of distance between released and not-released synaptic vesicles to the active zone at inhibitory synapses in Control (grey), Acute (orange) and Chronic (blue) epileptic mice. Paired t-test, Excitatory synapses: Control mice p = 0.0002 (n=40), Acute mice p = 0.0006 (n=41), Chronic mice p = 0.298 (n=112). Paired t-test, Inhibitory synapses: Control mice p = 0.06 (n=14), Acute mice p = 0.135 (n=13), Chronic mice p = 0.001 (n=28). **C**) 2D histograms of released vesicles distribution at excitatory (top) and inhibitory (bottom) synapses across the three conditions with active zone at the origin of the XY plane. Control (grey), Acute (orange) and Chronic (blue). Each synapse was spatially normalised (X and Y axes) and frequency is plotted on the Z axis. Scale bars: 0.1 normalised size X and Y; 0.1 fraction Z axis.

### Upregulation of synaptic proteins involved in vesicle positioning in Acute and Chronic epilepsy phases

To better understand molecular changes taking place in epileptic synapses, we performed an in-depth proteomic analysis of visual cortex synaptosomes. The expression profile of 1991 synaptic proteins extracted from animals in the acute and chronic phase of epilepsy was compared with controls. Using a fold change cut-off of 0.6, we found a total of 70 regulated proteins (51 proteins upregulated and 19 downregulated; **Fig. 3A, 3B**). As expected following TeNT injection, the Acute group showed a significant downregulation of VAMP1 and VAMP2 (Mainardi et al., 2012; Vannini et al., 2016). Interestingly, a few synaptic proteins remained upregulated at both stages of epilepsy, suggesting that one single TeNT injection is sufficient to induce persistent plastic changes. Proteins involved in synthesis of regulatory peptides, WNT pathway, immune response and membrane-trafficking were upregulated in hyperexcitable mice (i.e. Dickhopf related protein 3, Complement component 1q, Synaptotagmin 5, Semaphorin 4a, Carboxypeptidase E - CPE, Chromogranin B). The upregulation of neuropeptides was in line with previous reports (Vezzani and Sperk, 2004; Kovac and Walker, 2013; Clynen et al., 2014; Dobolyi et al., 2014; Nikitidou Ledri et al., 2016). These data prompted us to quantify the incidence of Dense Core Vesicles (DCV) in synaptic terminals. To this aim, we performed electron microscopy on samples collected from control and experimental animals and found no difference in the number of DCV across the three conditions (**Fig. 3C**). However, we found a tightening of synaptic vesicle clusters at excitatory synapses in both the Acute and Chronic groups and at inhibitory terminals in the Acute group (**Fig. 3D**). We limited our analysis to non-released vesicles, whose position is less likely to have been affected by recent recycling. Since the increase in CPE levels did not affect DCV incidence and given the changes in synaptic vesicles clustering at excitatory terminals in epileptic mice, we speculated that CPE might principally act through the pathways involved in vesicles organization (Ji et al., 2017).

**Figure 3:**
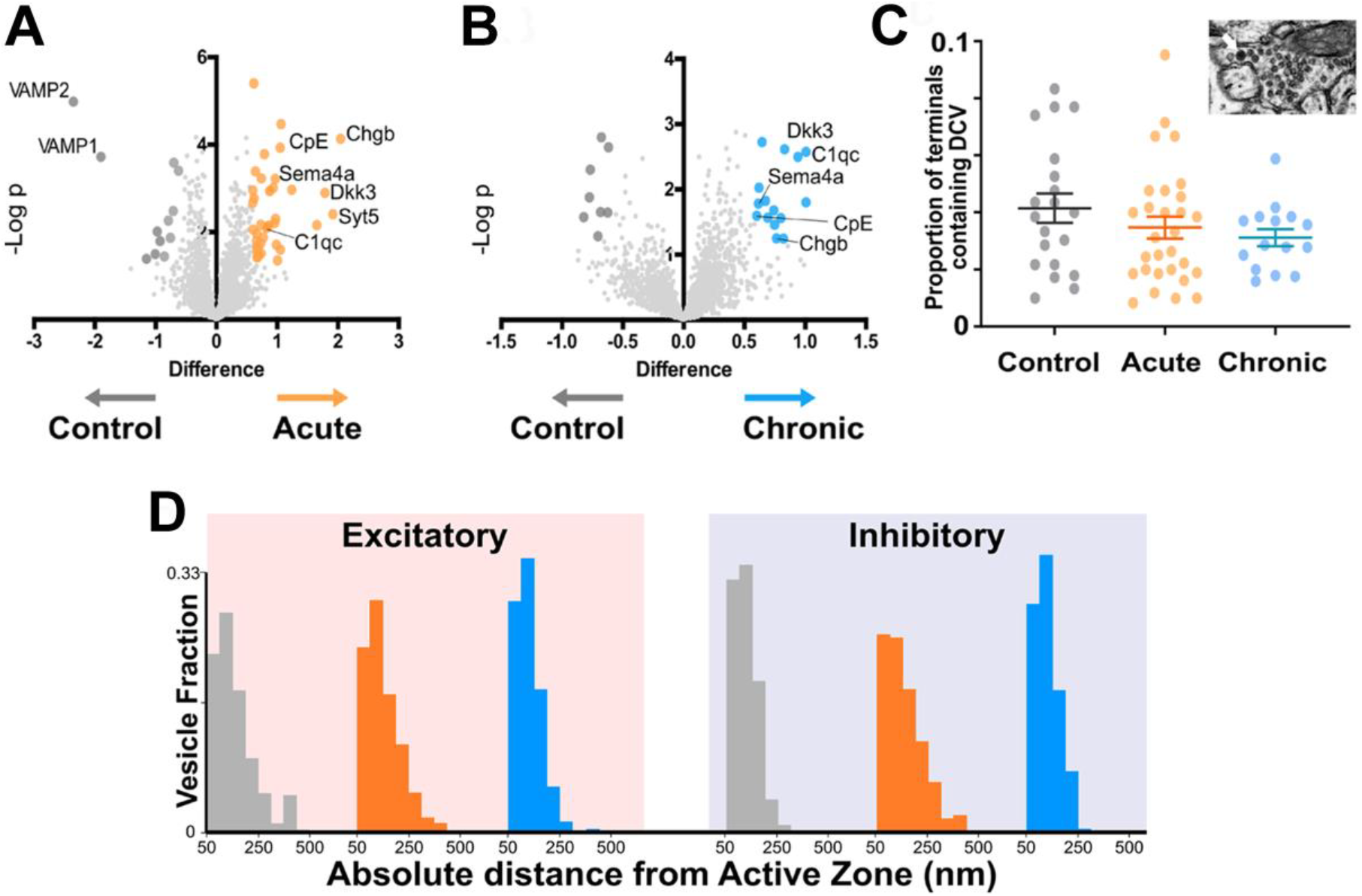
Proteomics analysis of synaptosomes reveal an increase of proteins involved in vesicular positioning. **A, B**) Differentially expressed proteins in Control vs Acute (**A)** and Chronic epileptic phase (**B**). Volcano plots are built plotting average ratio of TeNT vs. corresponding control against their t-test log P-values; significance thresholds: FDR > 0.05 and fold change > 0.6. Proteins significantly upregulated in Acute and Chronic tetanic animals are highlighted, respectively in orange and light blue; proteins significantly downregulated are in dark grey. Proteins abbreviations are Dkk3: Dickkopf-related protein 3; Sema4a: Semaphorin 4A; Cpe: carboxypeptidase e; Chgb: chromogranin b; Syt5: synaptotagmin5; VAMP1: Vesicle-associated membrane protein 1; VAMP2: Vesicle-associated membrane protein 2; C1qc: Complement C1q C Chain. **C)** Proportion of presynaptic terminals containing Dense Core Vesicles in different non-overlapping sampled areas of Control (grey; n=20), Acute (orange; n=29) and Chronic (blue; n=15) groups. No differences between groups (One Way ANOVA, p = 0.2869). Data are represented as mean ± SEM. Inset, a representative image of Dense Core Vesicles. **D)** Right: Distribution of distances of non-released vesicles from active zone at excitatory synapses in Control (grey; n=2140), Acute (orange; n=2503) and Chronic (blue; n=5705) groups (One-way ANOVA; F=238.15, p<0.0001, Control vs Acute: p<0.0001; Control vs Chronic: p<0.0001). Left: Distribution of distances of non-released vesicles from active zone at inhibitory synapses in Chronic (grey; n=543), Acute (orange; n=717) and Chronic (blue; n=1520) groups (F=75.57, p<0.0001, Control vs Acute: p<0.0001; Control vs Chronic: p>0.05).

### Acute Carboxypeptidase E inhibition reduces seizure activity in epileptic mice

Based on the indication that CPE is upregulated in TeNT-injected mice, and given its potential involvement in vesicle positioning (Ji et al., 2017), we decided to perform *in vivo* electrophysiological recordings in acute epileptic mice before and after pharmacological inhibition of CPE. We performed local field potential (LFP) recordings using a 16-channel silicon probe, spanning the whole cortical thickness in awake epileptic mice. Recording channels were divided in superficial (channels 1-5), intermediate (6-11) and deep (12-16) according to their position in the primary visual cortex. After baseline recording of seizures, we topically administered on the visual cortex GEMSA, a CPE inhibitor (**Fig. 4A, 4B**). The recording sessions following GEMSA administration showed a significant decrease in LFP coastline (**Fig. 4C**), indicating that CPE inhibition reduces indicators of seizure activity in epileptic mice.

**Figure 4:**
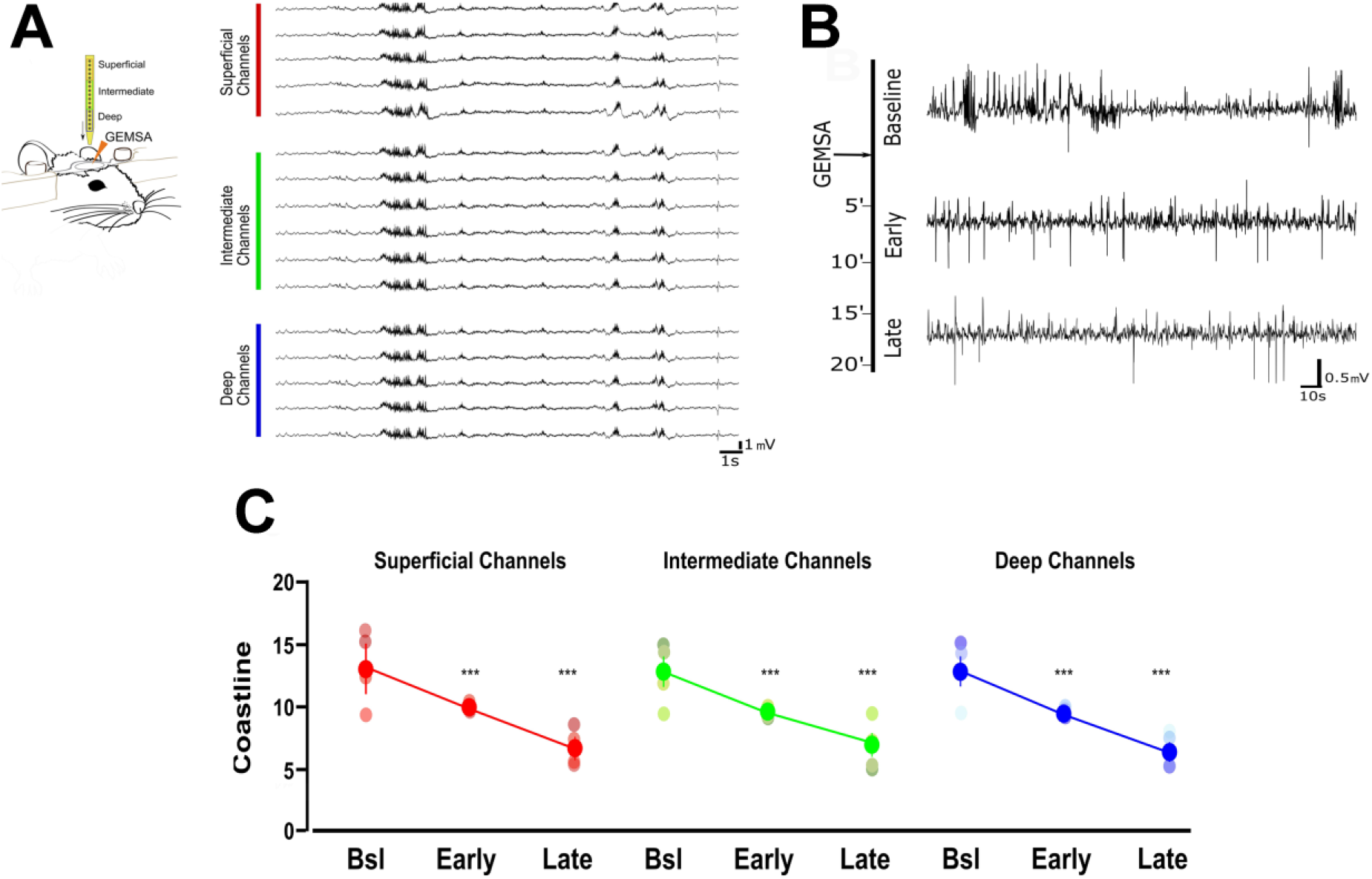
Acute inhibition of Carboxypeptidase (CPE) decreases hyperexcitability in TeNT-injected mice. **A**) Left: diagram of experimental design: a 16-channel silicone probe was used to record LFP in different layers of the primary visual cortex, channels were analysed in 3 groups according to their recording sites in relation to the surface of the cortex: the 5 most superficial, the 5 deepest and the 6 intermediate channels. GEMSA was applied locally to inhibit CPE activity. Right: Examples of LFP traces obtained with a 16-channels probe from the visual cortex of an Acute epileptic mouse. **B**) LFP traces of an Acute epileptic mouse before (baseline, top) and after GEMSA administration at two different time points: early (5 to 10 minutes) and late (10 to 20 minutes). **C**) Coastline analysis of LFP signals recorded before (baseline) and after GEMSA administration at early and late time points. The analysis was differentially performed for superficial (left, red), intermediate (middle, green) and deep (right, blue) channels (Two-way ANOVA, Channel factor p>0.05, Time factor p<0.001; Baseline vs Early: p < 0.01, Baseline vs Late: p < 0.001, Early vs Late: p < 0.001, n=4). The mean, SEM and value of individual recordings are shown for each group.

## Discussion

This study provides new insights into functional and ultrastructural synaptic changes in epileptic neuronal networks. Using a well-established model of epilepsy, we observed differential regulation of vesicular positioning and active zone size at excitatory and inhibitory synapses (**Fig. 1, 2**). We identified a homeostatic increase in active zone length specifically at inhibitory synapses, consistent with previous reports and our finding that GABA release is preferentially impaired by TeNT (Schiavo et al., 2000; Ferecskó et al., 2015). These early changes at inhibitory synapses are also consistent with previous observations made in the acute phase, when TeNT catalytic activity can still be detected (i.e. 10 days after TeNT-injection) (Mainardi et al., 2012; Vannini et al., 2016). We suggest that active zone length at inhibitory synapses is homeostatically upregulated in an attempt to restore baseline GABAergic release in spite of TeNT activity. We speculate that the lengthening of inhibitory active zones is later ‘discarded’, as a homeostatic mechanism, given its inability to overcome TeNT-induced reduction in vesicle release, as demonstrated by the reduced ratio of released GABAergic vesicles in both Acute and Chronic groups (**Fig. 1B**).

Ultrastructural changes of release competent vesicle positioning at excitatory synapses can only be detected at a later stage. In the chronic phase, excitatory terminals contained a smaller proportion of docked release-competent vesicles, consistent with the reported loss in spatial bias. A similar spatial reorganization of release competent vesicles can be achieved pharmacologically by stabilising actin, leading to a slower release rate during 10 Hz stimulation (Marra et al., 2012), its opposite, a tightening of synaptic vesicles, can be observed following Long-Term Potentiation in slices (Rey et al., 2020). Here, we report that spatial organization of release-competent synaptic vesicles can be modulated *in vivo*. During the chronic phase, released glutamatergic vesicles are positioned farther away from the active zone, potentially to limit their re-use during high-frequency activity. This loss of spatial bias may reduce the likelihood of generating spontaneous discharges in hyperexcitable networks. While a direct measure of the functional impact of this spatial reorganization is not possible with currently available methods, we can speculate that the reduction of released vesicles at the active zone of excitatory synapses may fit with the models of occupancy and two-step release proposed over the years by the Marty’s lab (Trigo et al., 2012; Pulido et al., 2015; Pulido and Marty, 2017; Miki et al., 2018). Interpreted in the light of Marty’s work, excitatory synapses in the chronic phase, although not changing in release fraction, may have a broader range of release latencies due to a reduction in occupancy at rest (Pulido et al., 2015; Pulido and Marty, 2017; Miki et al., 2018). Thus, in chronic epileptic mice the spatial organisation of release-competent vesicles farther from active zone may represent an attempt to homeostatically reduce networks’ synchronicity without affecting the total number of vesicles released. While not sufficient to block seizures in TeNT epileptic model, this spatial rearrangement may account for the reported reduction of seizures observed in the chronic phase (Vallone et al., 2016; Vannini et al., 2016; Chang et al., 2018). To dissect the molecular mechanisms underlying this change in spatial bias, we performed an unbiased analysis of synaptosomes content in the two different phases of epilepsy. Unsurprisingly, we found upregulation of several proteins involved in DCV trafficking as expected during intense synaptic remodelling. However, we did not find any statistical differences in the number of DCV present in the three experimental groups. We focussed our study on CPE, a protein that is involved in many different pathways, including neuropeptides’ synthesis and WNT/BDNF signalling, that was also hypothesized to regulate synaptic vesicles trafficking and positioning (Bamji et al., 2006; Staras et al., 2010; Skalka et al., 2016). Although the exact role of CPE is not completely clear, we reported a reduced hyperexcitability of acute epileptic mice after the administration of its inhibitor (**Fig. 4**). We also showed a tightening of synaptic vesicle clusters, measured from the active zone (**Fig. 3D, 3E**). Therefore our loss of spatial bias in recently released vesicles might happen on a background of overall contraction of vesicular clusters. This observation offers a possible interpretation for the effect of CPE inhibition on epileptiform activity, that was electrophysiologically measured *in vivo* (**Fig. 4**). The mechanisms by which CPE inhibition impacts on seizures remain to be fully clarified. CPE is involved in several different biosynthetic and signalling pathways (i.e. WNT, BDNF) which may account for the anti-epileptic effects. Moreover, CPE impact is likely to be indirect and very hard to dissect *in vivo*. However, further studies on other epileptic models are necessary to better elucidate CPE role in epilepsy. Taken together, our results suggest a complex landscape of molecular and ultrastructural changes evolving over time, opening intriguing questions regarding the temporal evolution of homeostatic changes in response to the induction of hyperexcitability. It would be particularly interesting to observe how homeostatic regulation of excitability adapts over development, for example in a model of genetic epilepsy, where epileptogenic factors are present since the very early formation of the nervous system (Lignani et al., 2020). Since TeNT-induced epilepsy is pharmacoresistant (Nilsen et al., 2005) and refractory seizures represent a major unmet medical need, drugs acting on CPE levels and other regulators of synaptic pools warrant further investigation as possible therapeutic treatments in currently intractable epilepsy.

## Acknowledgements

We thank Francesca Biondi (CNR Pisa) for the excellent animal care, Natalie Allcock and Ania Straatman-Iwanowska for the invaluable electron microscopy technical support and the scidraw.io team for the freely available vector graphics used in Figures 1 and 4. During manuscript elaboration, EV was supported by a postdoctoral fellowship released by Fondazione Umberto Veronesi (Milan, Italy). This work was funded by Epilepsy Research UK (pilot grant PGE1501 and project grant P1802) and by the Wellcome Trust Seed Award in Science (108201/Z/15/Z). We also gratefully acknowledge the financial support of CNR (Joint Laboratories project). This manuscript has been released as a pre-print at bioRxiv (Vannini et al., 2020) https://doi.org/10.1101/2020.01.05.895029.

## Author Contributions

E.V., M.C., V.M., L.MD conceived and designed the experiments; E.V., L.R., M.C. supplied the animal models; M.C., V.M. supervised the work; E.V., L.R., M.D., V.M. performed the experiments; E.V., M.D., V.M. analysed the data; E.V., V.M. wrote the paper. All authors contributed to the critical revision of the manuscript.

